# CLASP mediates microtubule repair by promoting tubulin incorporation into damaged lattices

**DOI:** 10.1101/809251

**Authors:** Amol Aher, Dipti Rai, Laura Schaedel, Jeremie Gaillard, Karin John, Laurent Blanchoin, Manuel Thery, Anna Akhmanova

**Author notes:** Lead contact: Anna Akhmanova.

## Abstract

Microtubule network plays a key role in cell division, motility and intracellular trafficking. Microtubule lattices are generally regarded as stable structures that undergo turnover through dynamic instability of their ends [1]. However, recent evidence suggests that microtubules also exchange tubulin dimers at the sites of lattice defects, which can either be induced by mechanical stress or occur spontaneously during polymerization [2–4]. Tubulin incorporation can restore microtubule integrity; moreover, “islands” of freshly incorporated GTP-tubulin can inhibit microtubule disassembly and promote rescues [3–7]. Microtubule repair occurs in vitro in the presence of tubulin alone [2–4, 8]. However, in cells, it is likely to be regulated by specific factors, the nature of which is currently unknown. CLASP is an interesting candidate for microtubule repair, because it induces microtubule nucleation, stimulates rescue and suppresses catastrophes by stabilizing incomplete growing plus ends with lagging protofilaments and promoting their conversion into complete ones [9–16]. Here, we used in vitro reconstitution assays combined with laser microsurgery and microfluidics to show that CLASP2α indeed stimulates microtubule lattice repair. CLASP2α promoted tubulin incorporation into damaged lattice sites thereby restoring microtubule integrity. Furthermore, it induced the formation of complete tubes from partial protofilament assemblies and inhibited microtubule softening caused by hydrodynamic flow-induced bending. A single CLASP2α domain, TOG2, which suppresses catastrophes when tethered to microtubules, was sufficient to stimulate microtubule repair, indicating that catastrophe suppression and lattice repair are mechanistically similar. Our results suggest that the cellular machinery controlling microtubule nucleation and growth can also help to maintain microtubule integrity.

## CLASP stalls depolymerization and promotes repair of microtubule lattices damaged by photoablation

To investigate whether CLASPs can promote microtubule repair, we modified previously described in vitro reconstitution assays with full length GFP-tagged CLASP2α purified from HEK293T cells (Figure S1A) [11]. Microtubules were grown from GMPCPP-stabilized seeds, visualized by adding fluorescently labeled tubulin and observed by Total Internal Reflection Fluorescence (TIRF) microscopy [11, 17]. In this assay, GFP-CLASP2α (Figure 1A) shows some binding to microtubule lattices and a weak enrichment at growing microtubule tips [11]. To explore the capacity of CLASP to repair microtubule lattices, we performed laser-mediated microsurgery experiments on dynamic microtubules. Lattice damage induced by laser irradiation next to a microtubule often led to progressive bending and eventual breakage of the microtubule at the damage site (Figures 1B,C and Video S1). This was most likely due to dissociation of tubulin dimers from the damaged microtubule lattice and subsequent loss of mechanical integrity. In the presence of tubulin alone, microtubules that bent by more than 10º after photo-damage typically broke (Figure 1C), although in 18% of the cases, microtubules straightened again, suggesting that they were repaired (Figure 1E). These data are in agreement with previous work showing that damaged microtubule lattices can be autonomously repaired by tubulin incorporation [2, 3, 8]. When these experiments were performed in the presence of CLASP2α, the protein rapidly accumulated at the site of damage (Figures 1D,F and Video S1), and the percentage of successful repair of microtubules bent at a damaged site by an angle of more than 10º increased threefold to 62% (Figure 1E).

**Figure 1.**
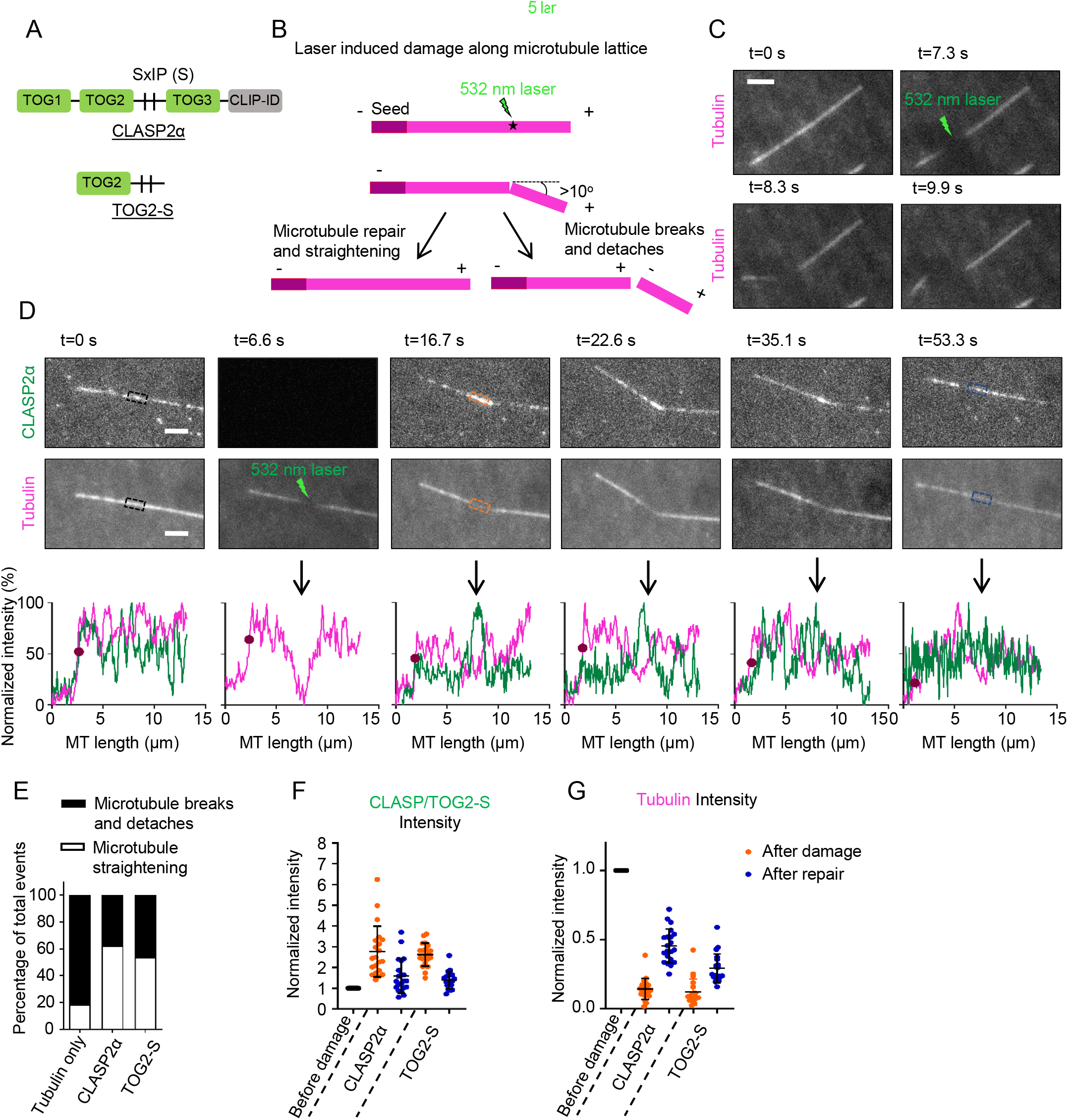
CLASP promotes repair of microtubule lattices damaged by laser illumination. (A) A scheme of full length CLASP2α and its TOG2-S fragment. Vertical lines labeled SxIP (Ser-any amino acid-Ile-Pro) represent EB-binding motifs located in the unstructured positively charged region adjacent to the TOG2 domain. (B) A scheme of the photodamage experiment using a 532 nm pulsed laser. Events when a microtubule bends to an angle >10^°^ at the point of photodamage result either in straightening of the lattice or microtubule breakage. (C,D) Stills from time lapse movies showing dynamic microtubules grown in the presence of Rhodamine-tubulin alone (C) or together with 30 nM GFP-CLASP2α (D), damaged by a pulsed 532 nm laser at a point along the lattice as indicated. In (D), intensity profiles along the microtubule for the CLASP (green) and tubulin channel (magenta) at different time points are shown in the bottom panels, with the arrow pointing to the site of photodamage. The purple circle on the plot indicates the end of the microtubule. Scale bars, 2 µm. (E) Plot showing the percentage of total events resulting in either microtubule breakage or straightening at the point of photo-damage in the presence of tubulin alone (n=22 microtubules analyzed from 4 experiments) or together with either 30 nM GFP-CLASP2α (n=53 microtubule analyzed from 6 experiments) or 30 nM GFP-TOG2-S (n=54 from 8 experiments). (F,G) Normalized intensity at the site of photodamage on the microtubule for the GFP channel for CLASP2α and TOG2-S (F) and Rhodamine-tubulin in the presence of CLASP2α and GFP-TOG2-S (G) before damage (black), immediately after damage (orange) and after repair (blue). n=21 microtubules from 4 experiments analyzed for CLASP2α, n=20 microtubules from 6 experiments for TOG2-S. See also Supplementary Figure S1 and Video S1.

Mammalian CLASPs contain three TOG-like domains, TOG1, TOG2 and TOG3, connected by flexible positively charged linkers, and a C-terminal domain (CLIP-ID) that binds to different partners and targets CLASPs to various subcellular locations [16, 18] (Figure 1A). Our previous work has shown that an isolated TOG2 domain has a very low affinity for microtubules and does not bind to free tubulin [11]. However, when TOG2 was fused to the adjacent intrinsically disordered positively charged protein region (a fusion protein termed TOG2-S, Figure 1A), TOG2 could bind to microtubule lattice, show some autonomous enrichment at growing microtubule ends and suppress catastrophes even in the absence of End Binding (EB) proteins, which normally target CLASPs to growing microtubule plus ends [11]. By performing laser damage experiments in the presence of TOG2-S, we found that it could also autonomously enrich at the damaged lattice site and promote microtubule repair (Figures 1E,F and S1A,B, Video S1). Importantly, measurements of the tubulin intensity at the damaged lattice site within microtubules grown in the presence of CLASP2α or TOG2-S showed that restoration of microtubule integrity was accompanied by incorporation of fresh tubulin (Figure 1G). These data show that CLASP2α and its TOG2-S fragment can autonomously recognize damaged microtubule lattices and enhance their restoration through tubulin incorporation.

Next, we set out to test if CLASP2α has a stabilizing effect on the plus end generated upon complete severing of a microtubule lattice. As was described previously [19], we found that severed microtubule lattices usually depolymerize from their freshly generated plus ends to the seed (81% of the microtubules), whereas the remaining 19% exhibited rescue along the lattice (Figures 2A,B,E and Video S2). In the presence of CLASP2α, microtubule plus-end depolymerization of newly generated plus ends was strongly inhibited: 53% of the microtubules promptly re-grew from the site of ablation (Figures 2A,C,E and Video S2). The remaining 47% were rescued along the dynamic lattice, in line with the previous findings that CLASP2α behaves as a rescue factor in our assays [11] (Figure S2A). TOG2-S fusion was also sufficient to suppress depolymerization of new plus ends generated by microtubule severing and promoted re-growth at the site of photo-ablation in 29% of the cases, although the protection was less efficient than with the full length CLASP2α (Figures 2D,E and S2B). Importantly, in the presence of either the full length CLASP2α or TOG2-S, none of the ablated microtubules depolymerized to the seed (Figure 2E), and most of these microtubules exhibited only very short depolymerization excursions (less than 1 µm) compared to tubulin alone (Figures 2F and S2A, B).

**Figure 2.**
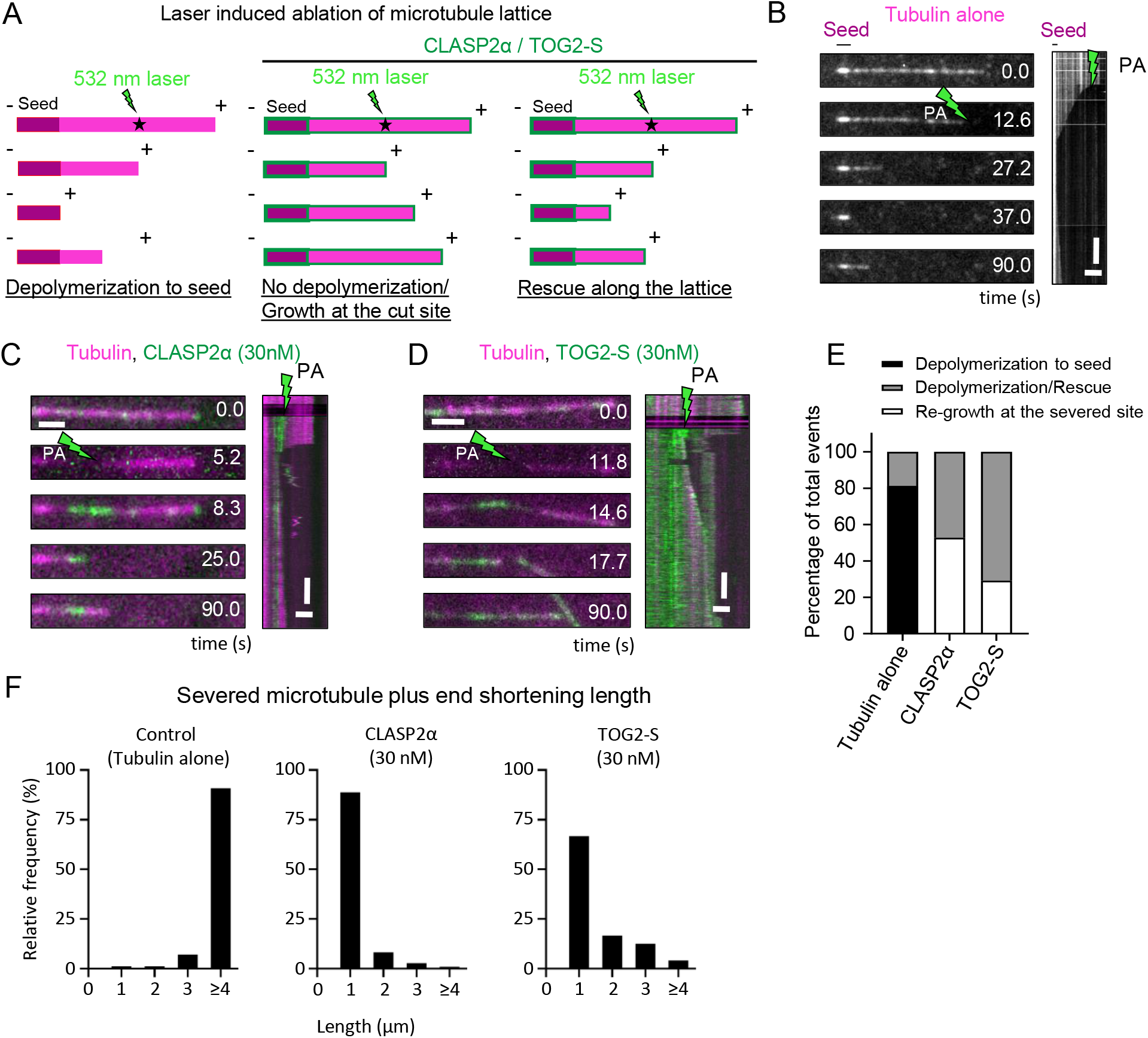
CLASP prevents depolymerization of laser-severed microtubules. (A) A scheme of the outcomes of laser severing experiments using a 532 nm pulsed laser in the presence of Rhodamine-tubulin alone or together with CLASP2α or TOG2-S. (B) Stills and the corresponding kymograph of a microtubule grown in the presence of Rhodamine-tubulin ablated with a 532 nm laser as indicated. Scale bars: still image, 2 µm; kymograph, 4 µm (horizontal) and 10 sec (vertical). (C,D) Stills and corresponding kymographs of microtubules grown in the presence of Rhodamine-tubulin together with either GFP-CLASP2α (30 nM) (C) or GFP-TOG2-S (30 nM) (D) ablated with a 532 nm laser as indicated (PA, photoablation). Scale bars as in panel (B). (E) Plot showing the percentage of total laser severing events resulting in either immediate microtubule regrowth at the site of photoablation, microtubule depolymerization to the seed or depolymerization followed by rescue along the lattice, in the presence of Rhodamine-tubulin alone or together with either 30 nM GFP-CLASP2α or 30 nM TOG2-S. (F) Plots showing the relative frequencies of microtubule plus end shortening lengths after laser-mediated severing of dynamic microtubules grown in the presence of Rhodamine-tubulin alone or together with either 30 nM GFP-CLASP2α or with 30 nM TOG2-S. For E and F, n=186 microtubules analyzed from 3 experiments for tubulin alone, n=36 microtubules analyzed from 3 experiments for CLASP2α and n=48 microtubules analyzed from 8 experiments for TOG2-S. See also Supplementary Figure S2 and Video S2.

## CLASP converts partial protofilament assemblies into complete tubes

The experiments described above suggest that CLASP promotes formation of a complete tube from a microtubule lacking some parts of protofilaments in the middle of the lattice. To test if CLASP has any effect on partial protofilament assemblies, we used a previously described engineered kinesin-5 dimer, which consists of the motor domain and neck linker of *Xenopus* kinesin-5 (Eg5) fused to the motor-proximal coiled coil derived from *Drosophila* kinesin-1 [20]. This engineered kinesin-5 was shown to enhance microtubule polymerization, likely by stabilizing a straight-tubulin conformation and enhancing lateral contacts between tubulin dimers [21]; it could also generate long tubulin ribbons and protofilament sheets at microtubule plus ends [20]. Similar curved microtubule plus-end extensions were also formed when CLASP2α or TOG2-S were added together with the Kin-5 dimer to the assay (Figures 3A-C, S1A, S3A,B and Video S3). However, whereas in the presence of the engineered kinesin-5 alone, these structures were transient and typically depolymerized, the addition of CLASP2α or TOG2-S to the assay led to generation of multiple straight microtubules from a single curled end (Figures 3C,D and S3A,B). These data suggest that CLASP2α and its TOG2-S fragment can each autonomously promote formation of complete microtubules from protofilament sheets. Since multiple microtubules can form from a single curled microtubule end most likely with a partial subset of protofilaments, these results suggest that in combination with kinesin-5 dimer, CLASP2α and TOG2-S not only help to repair strongly tapered microtubule ends, as it occurs during catastrophe suppression [11], but also likely promote extension of protofilament bundles from the side, allowing them to close into complete tubes. Given that a similar activity was displayed by TOG2-S, which, based on the competition with EB3 for microtubule tip accumulation, might be binding between protofilaments [11], it is possible that CLASP-mediated lattice repair activity depends on stabilizing lateral tubulin contacts in a microtubule.

**Figure 3.**
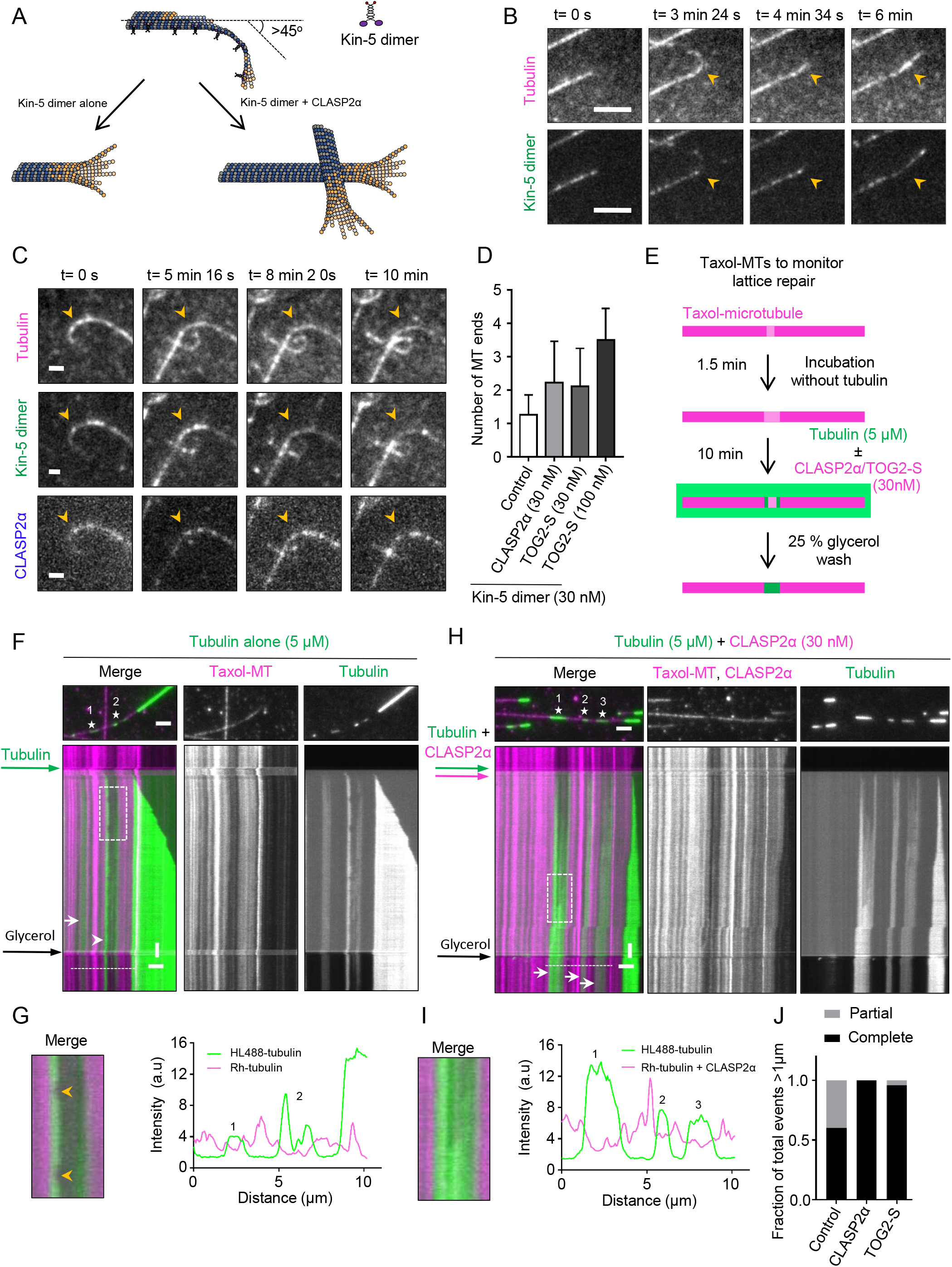
CLASP promotes formation of complete microtubules from partial protofilament assemblies. (A) A scheme of an experiment where tubulin sheet-or ribbon-like structures at the microtubule plus ends of dynamic microtubules are generated in the presence of Kinesin-5 dimer (Kin-5). (B,C) Stills from a time lapse showing a plus end of a microtubule grown in the presence of Rhodamine-tubulin and 30 nM Kinesin-5-GFP dimer (Kin-5) (B), or in the presence of Rhodamine-tubulin, 30 nM Kin-5-GFP and 30 nM TagBFP-CLASP2α (C). Scale bar: 2 µm. See also Video S3. (D) Plot showing the number of newly generated microtubule ends from a single microtubule plus end for microtubules grown in the presence of Rhodamine-tubulin and 30 nM Kin-5-GFP alone (n=38 microtubule plus ends analyzed from 3 experiments) or together with 30 nM TagBFP-CLASP2α (n=95 microtubule plus ends analyzed from 3 experiments), or with 30 nM TagBFP-TOG2-S (n=85 microtubule plus ends analyzed from 3 experiments), or with 100 nM Tag-BFP-TOG2-S (n=26 microtubule plus ends analyzed from 2 experiments). Events where the microtubule plus ends bent by angles over 45° with respect to the lattice were monitored in a 10 min time lapse. (E) Schematic representation of an experiment to monitor tubulin incorporation into the damaged microtubule lattices. Microtubule seeds prepared from Rhodamine-tubulin in the presence of Taxol were incubated in the wash buffer without Taxol and tubulin for 1.5 min to induce and expand lattice defects. Subsequently, they were incubated with 5 μM HiLyte Fluor 488-labeled tubulin with or without 30 nM mCherry-CLASP2α or 30 nM mCherry-TOG2-S. After 10 min, the residual free green tubulin was washed out with the wash buffer supplemented with 25% glycerol to prevent microtubule depolymerization, in order to better visualize incorporation of green tubulin into the damaged microtubule lattices. (F-I) Microtubule repair in the presence of tubulin alone (F, G) or together with 30 nM mCherry-CLASP2α (H, I). (F,H) Single frames (top) of time-lapse movies after the final washout and kymographs (bottom) showing green tubulin incorporation sites (numbered asterisks in stills) into Rhodamine-labeled microtubule lattices (magenta). In kymographs, white arrows indicate complete repair and white arrowheads partial repair. (G,I) Left, enlarged views of the boxed regions in the kymographs in panels F and H, respectively, showing partial or complete microtubule repair. Yellow arrowheads in panel G indicate events of loss of freshly incorporated tubulin. Right, intensity profiles along the microtubule for the Rhodamine-labeled microtubule seed channel (magenta) with or without mCherry-CLASP2α, and for the green tubulin channel. The numbers indicate incorporation sites specified in stills in panels F and H. Scale bars: 2 µm (horizontal) and 60 s (vertical). See also Video S4. (J) Plot showing the fraction of total events resulting in either complete or partial repair at the site of tubulin incorporations with the length exceeding 1 µm, in the presence of tubulin alone (n=15, analyzed from 2 experiments), together with either 30 nM mCherry-CLASP2α (n=22, analyzed from 5 experiments) or 30 nM mCherry-TOG2-S (n=23, analyzed from 4 experiments), where n is the total number of incorporations longer than 1 µm. See also Supplementary Figure S3.

## CLASP promotes complete repair of damaged microtubule lattices

To generate a substrate in which we could directly observe tubulin incorporation at the sites of lattice damage, we prepared Rhodamine-labeled microtubule seeds stabilized with the microtubule-stabilizing drug Taxol. Microtubules polymerized in the presence of Taxol are known to exhibit extensive lattice defects [22]; this property has been used, for example, to demonstrate the impact of microtubule defects on kinesin-based transport [23, 24]. Taxol-stabilized microtubules display more structural defects in the lattice when incubated with very low tubulin concentrations [25]. To promote defect formation, we incubated Taxol-stabilized microtubule seeds for 1.5 min in a buffer without Taxol and tubulin (Figure 3E). In these conditions, microtubule seeds undergo “erosion” by gradually loosing tubulin dimers from discrete sites, which can be detected as gaps with a reduced intensity of tubulin signal (Figures 3F-I, Video S4). When 5 µM tubulin with a green (HiLyte Fluor488) fluorescent label was added to such partially eroded Rhodamine-labeled microtubules, we observed that not only the seed ends were extended but also green tubulin was incorporated into the Rhodamine-labeled seed regions where the original tubulin signal was reduced (Figures 3F-I; Video S4). The addition of CLASP2α had no effect on the frequency of tubulin incorporation sites, as it depended on the number of Taxol-induced defects and the extent of seed erosion (Figure S3D). However, CLASP2α increased the percentage of damaged sites that appeared completely “healed” (Figures 3J, Video S4). This was because the polymerization of freshly added tubulin within the damaged sites was more processive, whereas in the presence of tubulin alone, incorporation was often confined to the edges of the gaps (Figure 3G, I). As a result, the average length of the analyzed incorporation sites was slightly longer in the presence of CLASP2α (Figure S3E). Moreover, in the absence of CLASP2α, the longest defects tended to break and were not included in the analysis. TOG2-S was equally efficient in this assay compared to full-length CLASP2α, indicating that microtubule-tethered activity of this domain alone is sufficient to promote repair (Figures 3J and S3C-E). Importantly, in this assay, free tubulin concentration was lower than in the assays described above (5 µM vs 15 µM in Figures 1, 2 and 3A-D), and therefore, we did not detect any incorporation of freshly added green tubulin signal along the lattice beyond the eroded zone, indicating that we were observing microtubule repair rather than initiation of freely growing microtubule protofilaments from the edges of the gaps within microtubule seeds. At higher tubulin concentrations in the presence of CLASP2α, we did sometimes observe protofilament overgrowth, which greatly complicated data interpretation, and this is why such conditions were not included in our analysis. Altogether, our results indicate that CLASP2α or its microtubule-tethered TOG2 domain promote processive tubulin addition to the ends of partial protofilament assemblies, allowing efficient repair of gaps in microtubule lattices.

## Microtubules exhibit increased resistance to mechanical stress in the presence of CLASP

Finally, we investigated the effect of CLASP on dynamic microtubules damaged in a more natural way, by inducing mechanical stress with a microfluidics setup described previously [2]. In order to study the impact of CLASP on the resistance of microtubules to mechanical forces, we applied cycles of hydrodynamic bending stress to microtubules in the absence or presence of CLASP2α. To this end, we grew dynamic microtubules in the absence of CLASP2α from Taxol-stabilized seeds attached to micropatterns inside a microfluidic device (Figure 4A). Microtubules were then bent by an orthogonal fluid flow for ten seconds either in the absence or in the presence of CLASP2α. The flow was subsequently stopped for ten seconds and the bending cycle was repeated (Figure 4B) [26]. Previous work showed that microtubules bend and subsequently straighten after each flow application, but the degree of softening (quantified as persistence length of microtubules) increases with each cycle due to the gradual loss of tubulin dimers from the microtubule lattice [2]. In the presence of tubulin alone, microtubules on average became 37% softer after six bending cycles, with 70% of the microtubules showing softening (Figure 4C,D and S4A). In the presence of CLASP2α, the average drop in the persistence length was much less pronounced (10%), with only 15% of the microtubules showing softening (Figure 4C,D and S4B). This indicates that microtubules exhibit increased resistance to mechanical stress in the presence of CLASP2α. These data support the idea that CLASP promotes repair of microtubule lattice damaged by bending and thus prevents progressive loss of tubulin dimers from bent lattice regions so that microtubules remain rigid.

**Figure 4.**
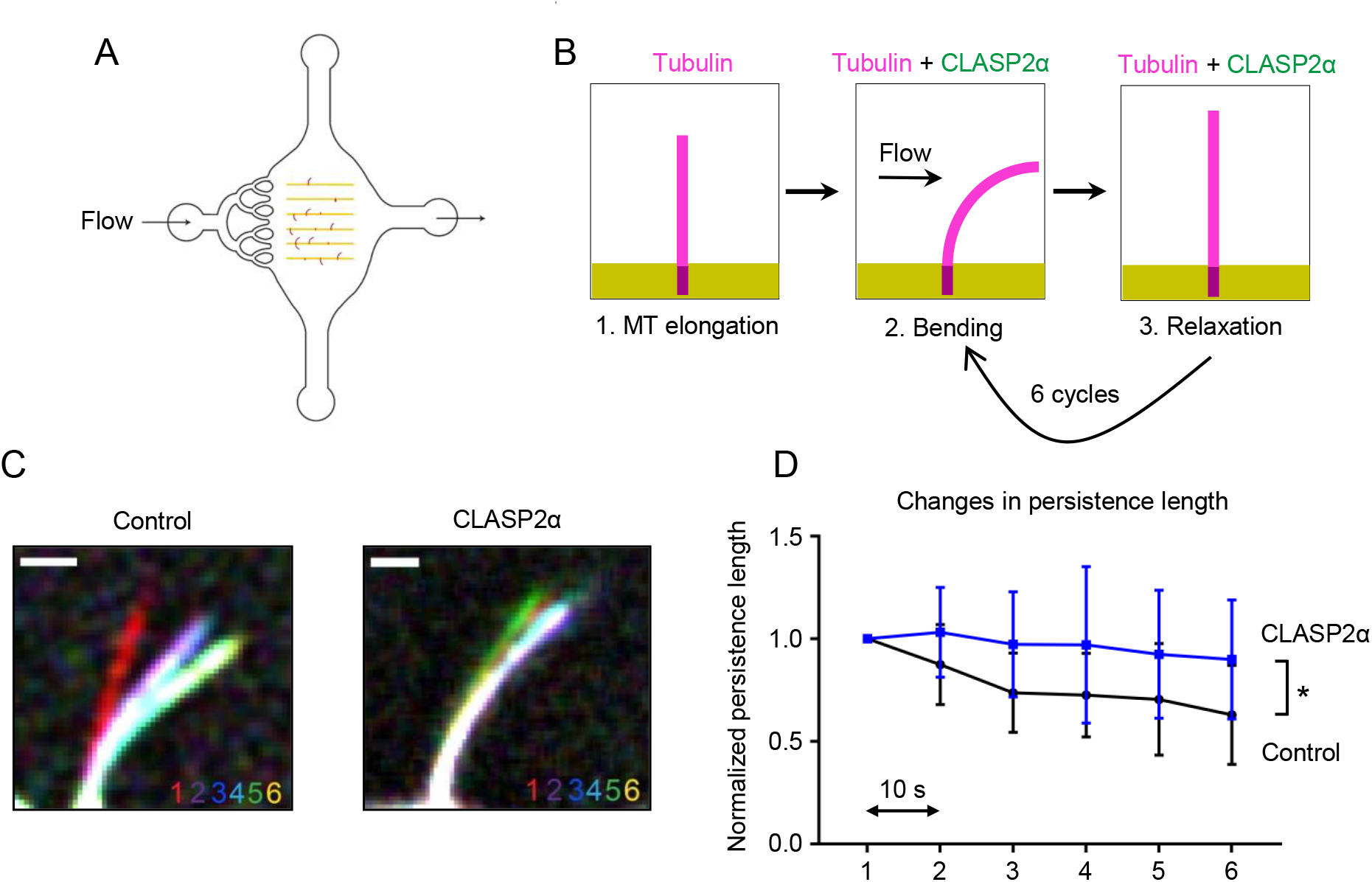
CLASP2α inhibits microtubule softening induced by hydrodynamic flow. (A) Illustration of the microfluidic device used for microtubule bending. (B) Scheme of the work sequence: 1. Red fluorescent microtubules were grown from seeds grafted on micropatterned lines. 2. Microtubules were bent for ten seconds by applying a fluid flow using the same mix as in 1, with or without 30 nM GFP-CLASP2α. 3. The flow was stopped for ten seconds before repeating the bending cycle. (C) The images show an overlay of maximum bent conformations where every cycle is represented in a different color. Scale bar: 3 µm. (D) Persistence length measured for microtubules bent in the presence (blue curve, n = 20) or absence (black curve, n = 23) of 30 nM GFP-CLASP2α. Persistence length was normalized to value in the first bending cycle for each microtubule. Values represent the average persistence length (mean ± SD) of individual measurements shown in Supplementary Figure S4. To test if microtubules showed softening, a Spearman correlation test for persistence length values over subsequent bending cycles was performed. It revealed significant softening of microtubules in both conditions (p = 0.01 and 0.08, respectively), though it was much less pronounced in the presence of CLASP2α. A t-test confirmed significant difference between the two curves (p = 0.006). See also Supplementary Figure S4.

## Conclusions

Taken together, our data reveal that CLASP2α is an autonomous microtubule repair factor. Since full length CLASP2α and its microtubule-tethered TOG2 domain were both active in microtubule repair assays, and since TOG2 is highly conserved between CLASP1 and CLASP2 and is present in all CLASP isoforms [18, 27], this property is likely shared by all mammalian CLASPs. Furthermore, given that previous work showed that the same TOG2-S fusion used here can suppress catastrophes [11], our data indicate that the mechanism of microtubule repair is similar to that of catastrophe suppression. Our previous work has shown that CLASPs can stabilize a microtubule plus end with an incomplete set of protofilaments, thereby promoting its recovery into a complete tube [11]. Furthermore, CLASPs are essential for microtubule nucleation from the Golgi in cells [28] and reduce the kinetic threshold for templated microtubule nucleation in vitro [11]. In the current study, we showed that CLASP2α promotes formation of complete tubes from tubulin sheets or ribbons generated by Kinesin-5 dimer, structures which are like microtubule nucleation intermediates [21]. All these activities likely depend on the ability of CLASPs to stabilize partial microtubule structures prone to depolymerization, shift the balance towards their polymerization and prevent their catastrophic disassembly. It is possible that the TOG2 domain of CLASP exerts this effect by stabilizing lateral contacts between tubulin dimers and thus enhances self-repair properties of microtubule lattices.

If CLASPs help to repair microtubules rapidly and efficiently, their loss could potentially lead to accumulation of microtubule damage that is repaired slowly. This might explain why CLASP depletion leads to increased GTP content and EB accumulation along microtubule shafts, a phenotype that could be rescued by the TOG2 domain of CLASP2 [29]. The strong reduction in the density of microtubule networks observed in CLASP-depleted cells could thus be caused not only by reduced microtubule nucleation and plus-end stability as assumed previously, but also by the decreased stability of microtubule lattices which are repaired less efficiently when CLASPs are absent. An important question for future research is whether and how other components of microtubule polymerization machinery can contribute to microtubule repair.

## Methods

### Protein purification for in vitro reconstitution assays

GFP-CLASP2α, GFP-TOG2-S, mCherry-CLASP2α, mCherry-TOG2-S, Tag-BFP-CLASP2α and Tag-BFP-TOG2-S used in the *in vitro* reconstitutions assays were purified from HEK293T cells using the Strep(II)-streptactin affinity purification as described previously (Figure S1A) [11]. Cells were transfected with the Strep-tagged constructs using polyethylenimine (PEI, Polysciences), in a ratio of 3:1 for PEI:DNA. Cells were harvested 2 days after transfection. Cells from a 15 cm dish were lysed in 500 µl of lysis buffer (50 mM HEPES, 300 mM NaCl and 0.5% Triton X-100, pH 7.4) supplemented with protease inhibitors (Roche) on ice for 15 minutes. The supernatant obtained from the cell lysate after centrifugation at 21,000 × g for 20 minutes was incubated with 40 µl of StrepTactin Sepharose beads (GE) pre-equilibrated with the lysis buffer for 45 minutes. The beads were washed 3 times in the lysis buffer without the protease inhibitors. The protein was eluted with 40 µl of elution buffer (50 mM HEPES, 150 mM NaCl, 1 mM MgCl_2_, 1 mM EGTA, 1 mM dithiothreitol (DTT), 2.5 mM d-Desthiobiotin and 0.05% Triton X-100, pH 7.4). Purified proteins were snap-frozen and stored at −80 °C. Kin-5-GFP was purified from *E. coli* BL-21 as described before (Figure S1A) [20].

### In vitro *microtubule dynamics assays*

Reconstitution of microtubule growth dynamics *in vitro* was performed as described previously [11]. GMPCPP-stabilized microtubule seeds (70% unlabeled tubulin, 18% biotin tubulin and 12% of Rhodamine-tubulin or HiLyte 488 tubulin) were prepared as described before [30]. Flow chambers, assembled from plasma-cleaned glass coverslips and microscopic slides were functionalized by sequential incubation with 0.2 mg/ml PLL-PEG-biotin (Susos AG, Switzerland) and 1 mg/ml NeutrAvidin (Invitrogen) in MRB80 buffer (80 mM piperazine-*N*,*N*[prime]-bis(2-ethanesulfonic acid (PIPES)), pH 6.8, supplemented with 4 mM MgCl_2_, and 1 mM EGTA). Microtubule seeds were attached to the coverslip through biotin-NeutrAvidin interactions. Flow chambers were further blocked with 1 mg/ml κ-casein. The reaction mixture with or without CLASP proteins (MRB80 buffer supplemented with 14.5 μM porcine brain tubulin, 0.5 μM Rhodamine-tubulin, 50 mM KCl, 1 mM guanosine triphosphate (GTP), 0.5 mg/ml κ-casein, 0.1% methylcellulose, and oxygen scavenger mix (50 mM glucose, 400 μg/ ml glucose oxidase, 200 μg/ml catalase, and 4 mM DTT)) was added to the flow chamber after centrifugation in an Airfuge for 5 minutes at 119,000 × *g*. For experiments with the Kin-5-GFP, KCL was excluded from the reaction mixture. The flow chamber was sealed with vacuum grease, and dynamic microtubules were imaged immediately at 30 °C using TIRF microscopy. All tubulin products were from Cytoskeleton Inc. for experiments in Figures 1–3.

### TIRF microscopy

*In vitro* reconstitution assays were imaged on a TIRF microscope setup as described previously [30] or on an iLas2 TIRF setup (see below). In brief, we used an inverted research microscope Nikon Eclipse Ti-E (Nikon) with the perfect focus system (Nikon), equipped with Nikon CFI Apo TIRF 100x 1.49 N.A. oil objective (Nikon) and controlled with MetaMorph 7.7.5 software (Molecular Devices). The microscope was equipped with TIRF-E motorized TIRF illuminator modified by Roper Scientific France/PICT-IBiSA, Institut Curie. To keep the *in vitro* samples at 30 °C, a stage top incubator model INUBG2E-ZILCS (Tokai Hit) was used. For excitation, 491 nm 100 mW Calypso (Cobolt) and 561 nm 100 mW Jive (Cobolt) lasers were used. We used ET-GFP 49002 filter set (Chroma) for imaging of proteins tagged with GFP or ET-mCherry 49008 filter set (Chroma) for imaging of proteins tagged with mCherry. Fluorescence was detected using an EMCCD Evolve 512 camera (Roper Scientific) with the intermediate lens 2.5X (Nikon C mount adapter 2.5X) or using the CoolSNAP HQ2 CCD camera (Roper Scientific) without an additional lens. In both cases the final magnification was 0.063 μm/pixel.

The iLas2 system (Roper Scientific) is a dual laser illuminator for azimuthal spinning TIRF (or Hilo) illumination and with a custom modification for targeted photomanipulation. This system was installed on Nikon Ti microscope (with the perfect focus system, Nikon), equipped with 150 mW 488 nm laser and 100 mW 561 nm laser, 49002 and 49008 Chroma filter sets, EMCCD Evolve mono FW DELTA 512×512 camera (Roper Scientific) with the intermediate lens 2.5X (Nikon C mount adapter 2.5X), CCD camera CoolSNAP MYO M-USB-14-AC (Roper Scientific) and controlled with MetaMorph 7.8.8 software (Molecular Device). To keep the *in vitro* samples at 30^°^C, a stage top incubator model INUBG2E-ZILCS (Tokai Hit) was used. The final resolution using EMCCD camera was 0.065 µm/pixel, using CCD camera it was 0.045 µm/pixel.

Both microscopes were equipped with an iLas system (Roper Scientific France/PICT-IBiSA) for FRAP and photoablation. The 532 nm Q-switched pulsed laser (Teem Photonics) was used for photoablation by targeting the laser on the microtubule lattice on the TIRF microscope and next to the lattice to induce damage.

### Microtubule repair assays with Taxol-stabilized microtubules

Taxol-stabilized microtubule seeds were prepared by polymerizing 29 μM porcine brain tubulin containing 13% biotinylated-tubulin and 6% Rhodamine-labeled tubulin in MRB80 buffer supplemented with 2 mM GTP at 37°C for 30 min. Taxol (Enzo Life Sciences) (18 µM) was then added to the tubulin-GTP mix and seeds were then sedimented by centrifugation at 16,200 ×g for 15 min at room temperature. Finally, the pellet was resuspended in warm 10 μM Taxol solution in MRB80 buffer. Taxol-stabilized microtubule seeds were then wrapped with aluminium foil and stored at room temperature for a maximum of 2 weeks.

For tubulin incorporation experiments, Taxol-stabilized microtubule seeds were immobilized in the flow chamber and were washed immediately with the wash buffer (80 mM PIPES, 4 mM MgCl_2_, 1 mM EGTA, 50 mM KCl, 0.5 mg/ml κ-casein, 0.1% methylcellulose, and oxygen scavenger mix (50 mM glucose, 400 μg/ml glucose-oxidase, 200 μg/ml catalase, and 4 mM DTT)). Time-lapse movies were immediately started on the TIRF microscope at 30°C with a 2 s time interval and 100 ms exposure time for 25 minutes. During the imaging session, seeds were incubated in the wash buffer without Taxol and tubulin for 1.5 min to promote lattice defect formation. Subsequently, they were incubated in MRB80 buffer supplemented with 5 μM HiLyte Fluor 488-labeled tubulin, 50 mM KCl, 1 mM GTP, 0.5 mg/ml κ-casein, 0.1% methylcellulose, and oxygen scavenger mix (50 mM glucose, 400 μg/ml glucose-oxidase, 200 μg/ml catalase, and 4 mM DTT) with or without 30 nM mCherry-CLASP2α or 30 nM mCherry-TOG2-S for 10 min to promote repair. Finally, the residual free green tubulin was washed out with the wash buffer supplemented with 25% glycerol to prevent microtubule seed depolymerization and to clearly visualize incorporation of green tubulin into the damaged microtubule lattices.

### Microtubule repair assays with mechanically damaged microtubules

#### Tubulin purification and labeling

For microtubule bending experiments, tubulin was purified from fresh bovine brain by three cycles of temperature-dependent assembly and disassembly in Brinkley Buffer 80 (BRB80 buffer: 80 mM PIPES, pH 6.8, 1 mM EGTA, 1 mM MgCl_2_ plus 1 mM GTP). MAP-free brain tubulin was purified by cation-exchange chromatography (Fractogel EMD SO, 650 M, Merck) in 50 mM PIPES, pH 6.8, supplemented with 1 mM MgCl_2_ and 1 mM EGTA. Purified tubulin was obtained after a cycle of polymerization and depolymerization. Fluorescent tubulin (ATTO-565-labeled tubulin) and biotinylated tubulin were prepared as follows: Microtubules were polymerized from brain tubulin at 37°C for 30 min and layered onto cushions of 0.1 M Na-HEPES, pH 8.6, 1 mM MgCl_2_, 1 mM EGTA, 60% v/v glycerol, and sedimented by high-speed centrifugation at 30°C. Then, microtubules were resuspended in 0.1 M Na-HEPES, pH 8.6, 1 mM NHS-ATTO (ATTO Tec), or NHS-Biotin (Pierce) for 10 min at 37°C. The labeling reaction was stopped using 2 volumes of 2x BRB80, containing 100 mM potassium glutamate and 40% v/v glycerol, and then microtubules were sedimented onto cushions of BRB80 supplemented with 60% glycerol. Microtubules were resuspended in cold BRB80. Microtubules were then depolymerized and a second cycle of polymerization and depolymerization was performed before use.

#### Cover glass micropatterning

The micropatterning technique was adapted from [26]. Cover glasses were cleaned by successive chemical treatments: 30 min in acetone, 15 min in ethanol (96%), rinsing in ultrapure water, 2 h in Hellmanex III (2% in water, Hellmanex), and rinsing in ultrapure water. Cover glasses were dried using nitrogen gas flow and incubated for three days in a solution of tri-ethoxy-silane-PEG (30 kDa, PSB-2014, Creative PEGWorks) 1 mg/ml in ethanol (96%) and 0.02% HCl, with gentle agitation at room temperature. Cover glasses were then successively washed in ethanol and ultrapure water, dried with nitrogen gas, and stored at 4°C. Passivated cover glasses were placed into contact with a photomask (Toppan) with a custom-made vacuum-compatible holder and exposed to deep UV (7 mW/cm^2^ at 184 nm, Jelight) for 2.5 min. Deep UV exposure through the transparent micropatterns on the photomask created oxidized micropatterned regions on the PEG-coated cover glasses.

#### Microfluidic circuit fabrication and flow control

The microfluidic device was fabricated in polydimethylsiloxane (PDMS, Sylgard 184, Dow Corning) using standard photolithography and soft lithography. The master mold was fabricated by patterning 85 µm thick negative photoresist (SU8 3050, Microchem, MA) by photolithography. A positive replica was fabricated by replica molding PDMS against the master. Prior to molding, the master mold was silanized with trichloro(1H,1H,2H,2H-perfluorooctyl)silane (Sigma) for easier lift-off. Four inlet and outlet ports were made in the PDMS device using 0.5 mm soft substrate punches (UniCore 0.5, Ted Pella, Redding, CA). The PDMS device was then brought into contact with a micropatterned cover glass and placed in a custom-made holder that could be fitted on the microscope stage. A transparent plate was fixed on the holder to apply gentle pressure on the chip in order to avoid leaks without the need of permanent bonding to the cover glass. The top plate had four openings for the inlet and outlet tubing. Teflon tubing (Tefzel, inner diameter: 0.03’’, outer diameter: 1/16’’, Upchurch Scientific) was inserted into the two ports serving as outlets. Tubing with 0.01’’ inner and 1/16’’ outer diameter was used to connect the inlets via two three-way valves (Omnifit Labware, Cambridge, UK) that could be opened and closed by hand to a computer-controlled microfluidic pump (MFCS-4C, Fluigent, Villejuif, France). Flow inside the chip was controlled using the MFCS-Flex control software (Fluigent). Custom rubber pieces that fit onto the tubing were used to close the open ends of the outlet tubing when needed.

#### Microtubule growth on micropatterns

Microtubule seeds were prepared at 10 µM tubulin concentration (30% ATTO-565 or ATTO-488-labeled tubulin and 70% biotinylated tubulin) in BRB80 supplemented with 0.5 mM GMPCPP at 37°C for 1 h. The seeds were incubated with 1 µM Taxotere (Sigma) at room temperature for 30 min and were then sedimented by high speed centrifugation at 30°C and resuspended in BRB80 supplemented with 0.5 mM GMPCPP and 1 µM Taxotere. Seeds were stored in liquid nitrogen and quickly warmed to 37°C before use.

The holder with the chip was fixed on the stage and the chip was perfused with NeutrAvidin (25 µg/ml in BRB80, Pierce), then washed with BRB80, passivated for 20s with PLL-g-PEG (Pll 20K-G35-PEG2K, Jenkam Technology) at 0.1 mg/ml in 10 mM Na-HEPES (pH 7.4), and washed again with BRB80. Microtubule seeds were flown into the chamber at high flow rates perpendicular to the micropatterned lines to ensure proper orientation of the seeds. Unattached seeds were washed out immediately using BRB80 supplemented with 1% BSA. Seeds were elongated with a mixture containing 27 µM tubulin (20% labeled) in BRB80 supplemented with 50 mM NaCl, 25 mM NaPi, 1 mM GTP, an oxygen scavenger cocktail (20 mM DTT, 1.2 mg/ml glucose, 8 µg/ml catalase and 40 µg/ml glucose oxidase), 0.1% BSA and 0.033% methyl cellulose (1500 cp, Sigma). Microtubules were bent by an orthogonal fluid flow either using the same mixture supplemented with 0.02% red fluorescent beads (0.52 µm diameter, Thermo Scientific) or supplementing it additionally with 30 nM GFP-CLASP2α.

#### Imaging

Microtubules were visualized using an objective-based azimuthal ILAS2 TIRF microscope (Nikon Eclipse Ti, modified by Roper Scientific) and an Evolve 512 camera (Photometrics). The microscope stage was kept at 35°C using a warm stage controller (LINKAM MC60). Excitation was achieved using 491 and 561 nm lasers (Optical Insights). Time-lapse recording was performed using Metamorph software (version 7.7.5, Universal Imaging). Movies were processed to improve the signal/noise ratio (smooth and subtract background functions of ImageJ, version 2.2.0-rc-65 / 1.51s).

#### Measurement of microtubule persistence length

The microtubule is described as an inextensible slender filament with length L and bending rigidity κ, which is bent in two dimensions by the fluid flow. Its elastic energy E is given by

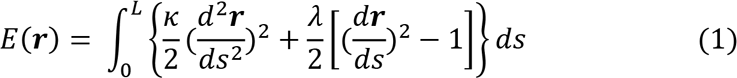

The vector **r**(s) denotes the position of the filament parameterized by the arc length s and λ denotes a Lagrange multiplier associated with the inextensibility condition |d**r**/ds|=1. The force exerted on the filament is given by the functional variation of the potential E with respect to the filament position vector **r**

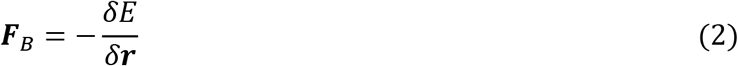

The filament orientation is fixed by the seed orientation at s=0, whereas the other end of the filament at s=L is force-free. The hydrodynamic drag exerted by the fluid flow on a slender filament is given by

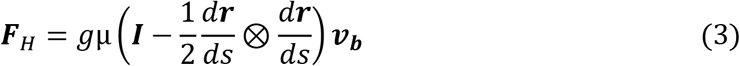

where **v**_b_ denotes the velocity field measured by the bead displacements, µ denotes the viscosity of the fluid and g denotes a geometrical factor of the order of 1, which depends on the distance of the filament from the surface, the radius of the filament and the distance of the beads from the surface. ⊗ denotes the outer product and *I* is the identity tensor. In mechanical equilibrium

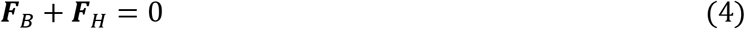

which determines the equilibrium shape of the filament subject to the appropriate boundary conditions. The filament rigidity was determined by solving equations (1)–(4) using the AUTO-07p software package and by minimizing the function

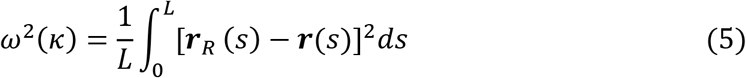

where **r**_R_(s) denotes the measured position of the filament. The persistence length is then given by Lp =κ/(k_B_ T). ω is a measure for the distance between the shapes of two microtubules. In the fitting routine for the experimentally measured microtubule shapes, ω denotes the distance between the shape of the experimental snake and an inextensible flexible filament subjected to the same flow as the experimental snake. We assumed that the origin of the microtubule was clamped in the direction of the seed. To correct for a measurement error of the microtubule origin, we optimized equation (5) also for the position of the microtubule origin.

## Author Contributions

A.Ah, D.R, L.S., M.T., L.B. and A.Ak. designed experiments and wrote the paper. A.Ah., D.R. L.S., J.G., K.J. performed experiments and data analysis, A.Ak. coordinated the project.

## Acknowledgements

We thank Dr. Kai Jiang, Wuhan University for suggesting experiments with Kin-5 dimer. This work was supported by the European Research Council Synergy grant 609822 to A.Ak. and European Research Council Consolidator grant 771599 to MT.

---

**Supplementary Figure S1.**
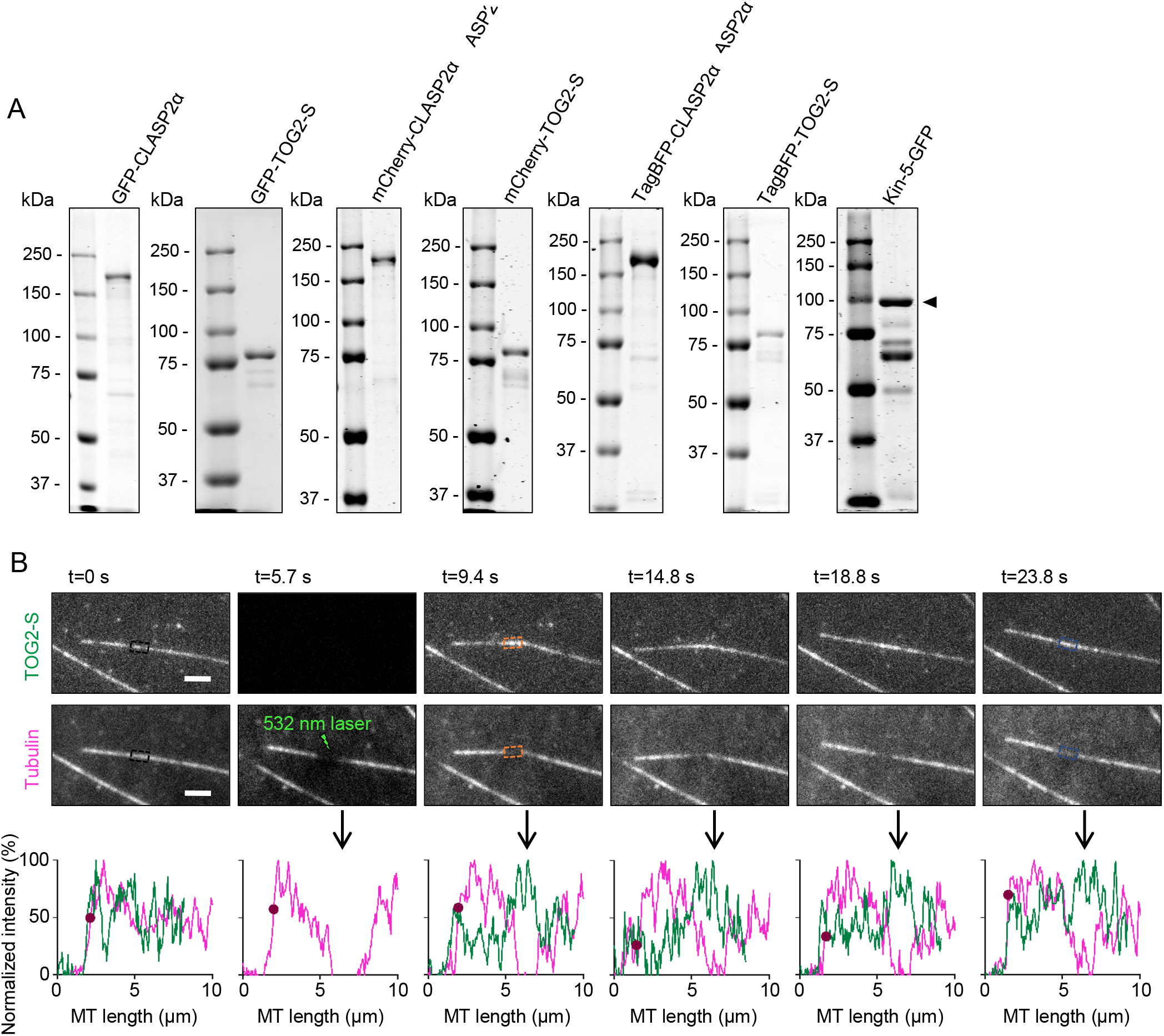
TOG2-S promotes repair of microtubule lattices damaged by laser illumination. (A) Proteins purified from HEK293T cells (CLASP2α and TOG2-S constructs) and *E.coli* (Kin-5-GFP) used in this study analyzed by SDS-PAGE. (B) Stills from a time lapse of a dynamic microtubule grown in the presence of 30 nM GFP-TOG2-S (upper panel) and Rhodamine-tubulin (lower panel) damaged at a point along the lattice as indicated. Intensity profiles along the microtubule for the TOG2-S and tubulin channel at different time points are shown below, with the arrow pointing to the site of photodamage. The purple circle on the plot indicates the end of the microtubule. Scale bar: 2 µm. See also Video S1.

**Supplementary Figure S2.**
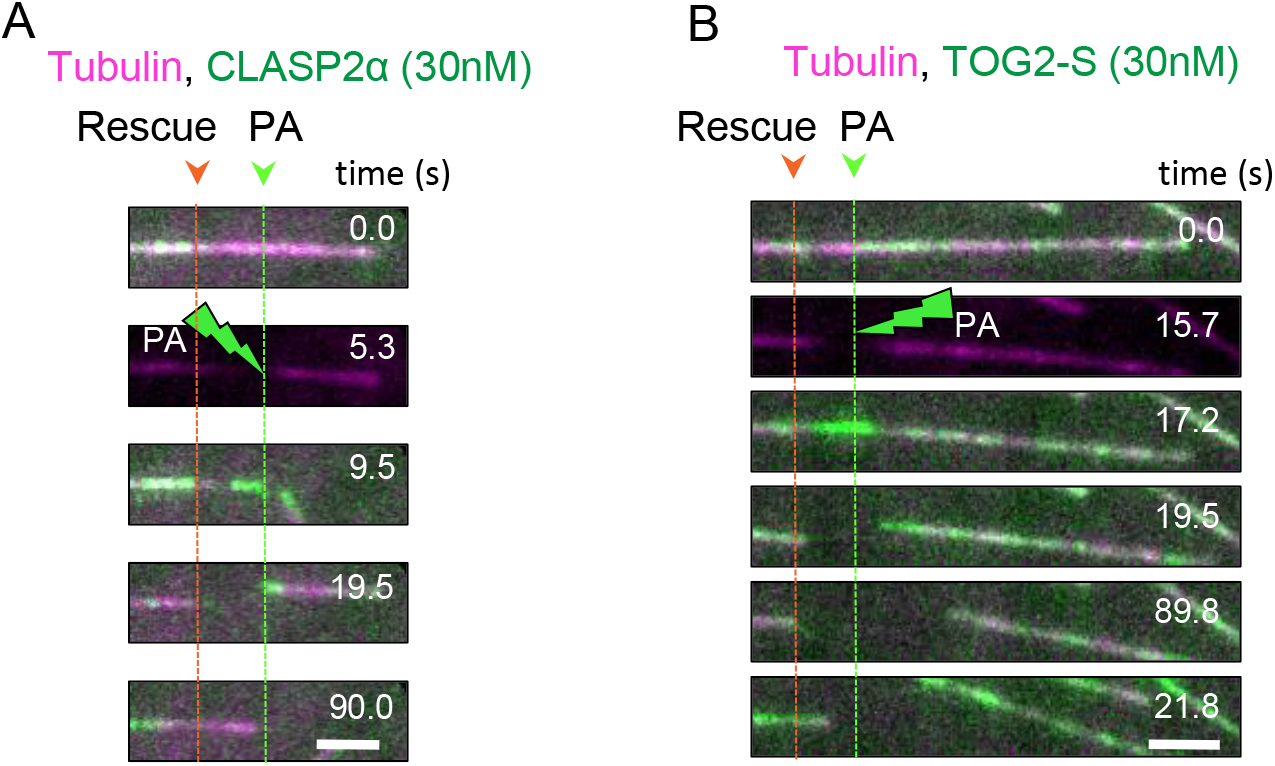
CLASP2α and TOG2-S induce rescue of laser severed microtubules. (A,B) Stills illustrating microtubule rescue after laser severing of microtubules grown in the presence of Rhodamine-tubulin together with either 30 nM GFP-CLASP2α (A) or 30 nM GFP-TOG2-S (B). Green arrowheads and dotted lines indicate the position of microtubule photoablation (PA) and orange arrowheads and dotted lines indicate the position of microtubule rescue. Scale bars: 2 µm.

**Supplementary Figure S3.**
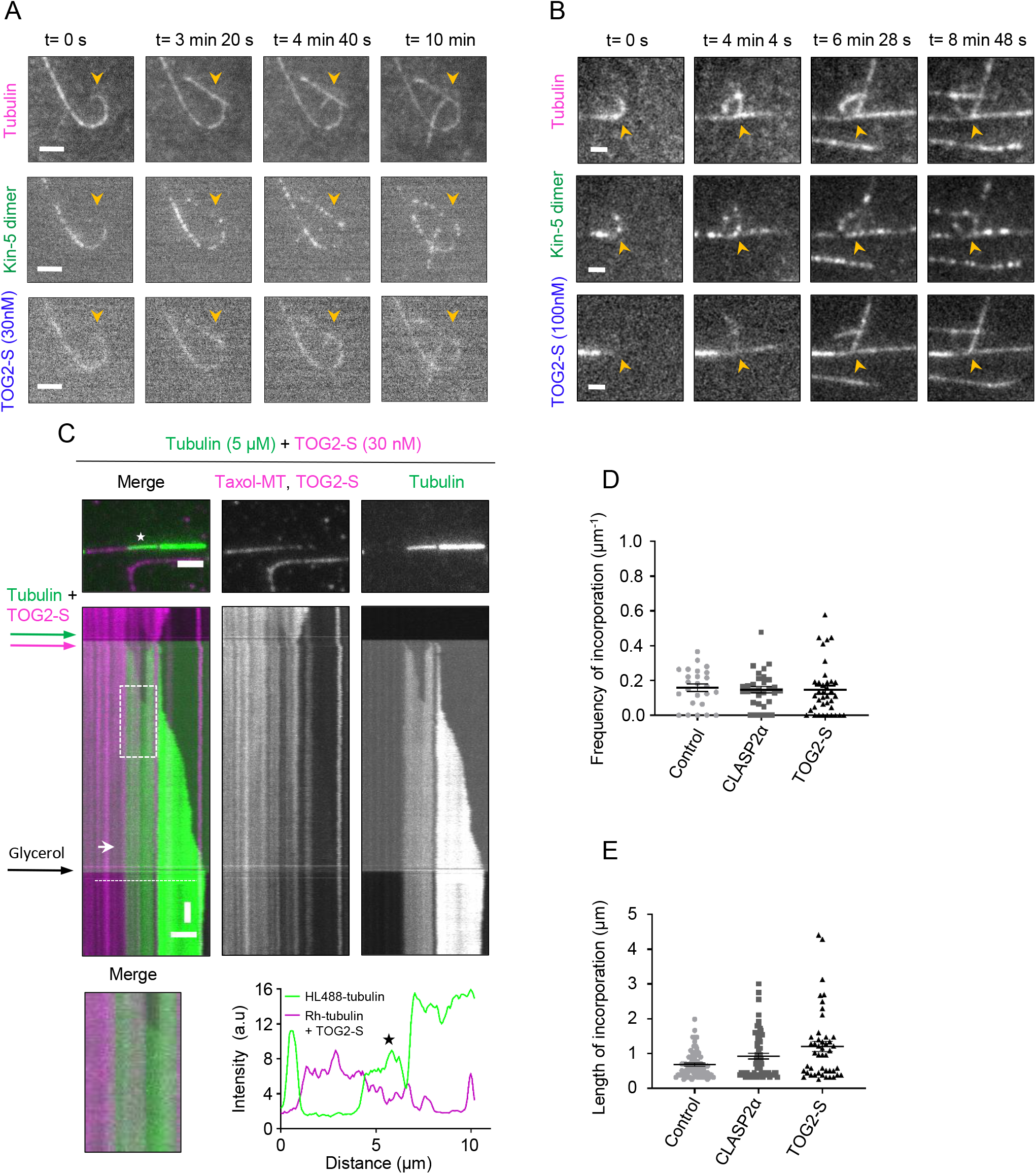
CLASP promotes formation of complete microtubules from partial protofilament assemblies. (A,B) Stills from a time lapse showing a microtubule plus end grown in the presence of Rhodamine-tubulin, 30 nM Kin-5-GFP together with either 30 nM TagBFP-TOG2-S (A) or 100 nM TagBFP-TOG2-S. Scale bar: 3 µm. See also Video S3. (C) Microtubule repair in the presence of tubulin together with 30 nM mCherry-TOG2-S. Single frames (top) of a time-lapse movie after the final washout and kymographs (middle) showing green tubulin incorporation spots (asterisk in the still image) into microtubule lattices (magenta). In the kymograph, white arrow indicates complete repair. Bottom left, enlarged view of the boxed area in the kymograph above, showing complete microtubule repair. Bottom right, an intensity profile along the microtubule for the Rhodamine-labeled microtubule seed channel and mCherry-TOG2-S (magenta) and for the green tubulin channel. The asterisk indicates the incorporation site specified in the still image above. Scale bars: 2 µm (horizontal) and 60 s (vertical). See also Video S4. (D,E) Frequency of incorporation per unit length per microtubule (D) and average length of incorporations (E) in the presence of tubulin alone (n=73, M=25, L=459.25 µm, analyzed from 2 experiments), together with 30 nM mCherry-CLASP2α (n=64, M=31, L=450.74 µm, analyzed from 5 experiments) or 30 nM mCherry-TOG2-S (n=52, M=37, L=418.43 µm, analyzed from 4 experiments), where n, M and L are total number of incorporations, total number of microtubules and total length of microtubules analyzed respectively. Error bars represent SEM.

**Supplementary Figure S4.**
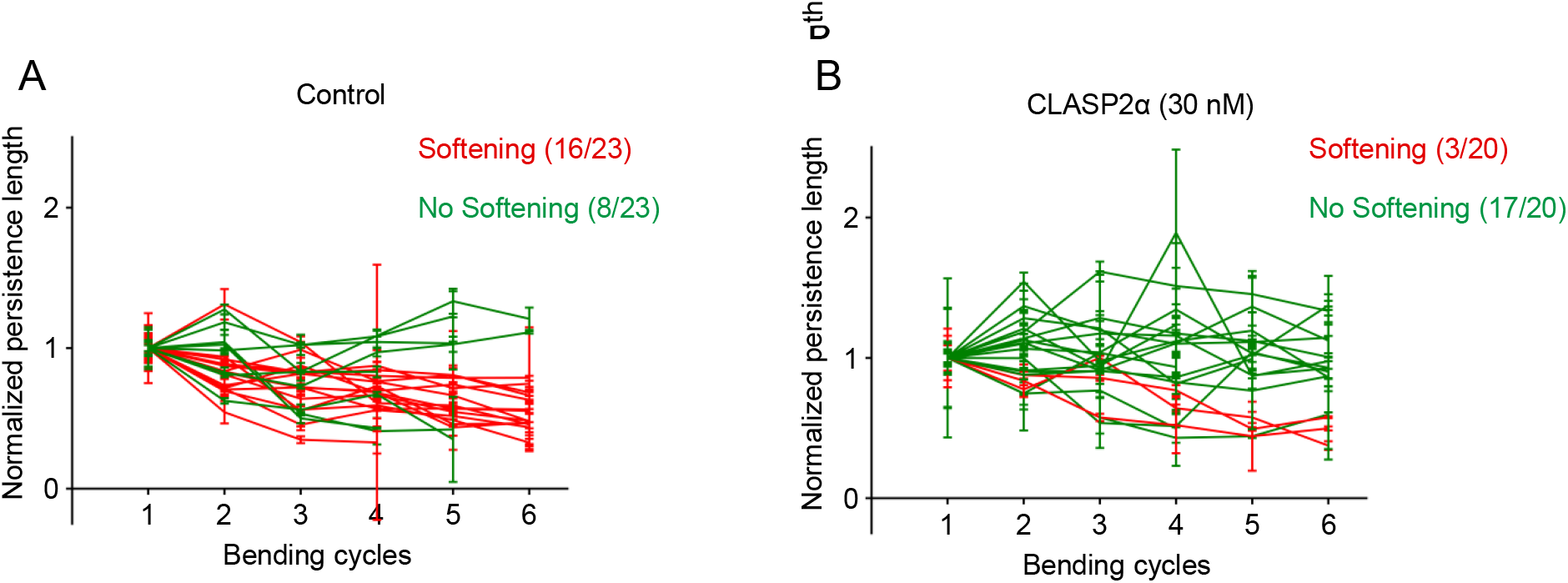
Persistence length measurements of microtubules bent in the absence or presence of CLASP2α. (A-B) Individual persistence length measurements of microtubules bent in the absence (A) and presence (B) of 30 nM GFP-CLASP2α. Values represent the average of five independent measurements for each bending cycle (mean ± SD) and were normalized to the initial value. A Spearman correlation test was performed to test for softening. Red lines indicate microtubules that became significantly softer and green lines indicate microtubules that did not show significant softening.

## Supplemental videos

**Video S1, related to Figure 1 and Supplementary Figure S1. CLASP promotes repair of microtubule lattices damaged by laser illumination.**

Microtubule lattice repair at the laser induced damage site in a microtubule grown in the presence of Rhodamine-tubulin alone (upper panel, corresponds to Figure 1C), a microtubule grown in the presence of Rhodamine-tubulin (magenta) and 30 nM GFP-CLASP2α (green) (middle panel, corresponds to Figure 1D) and a microtubule grown in the presence of Rhodamine-tubulin (magenta) and 30 nM GFP-TOG2-S (green) (lowermost panel, corresponds to Supplementary Figure S1). Yellow arrowheads point to the site of laser induced photodamage. Images were acquired using a TIRF microscope in a stream mode at a 100 ms interval. Video is sped up 50 times. Time is shown in seconds.

**Video S2, related to Figure 2. CLASP prevents depolymerization of laser-severed microtubules.**

Microtubules were ablated with a laser in the presence of Rhodamine-tubulin alone (upper panel, corresponds to Figure 2B), in the presence of Rhodamine-tubulin (magenta) and 30 nM GFP-CLASP2α (green) (middle panel, corresponds to Figure 2C) or in the presence of Rhodamine-tubulin (magenta) and 30 nM GFP-TOG2-S (green) (lowermost panel, corresponds to Figure 2D). Yellow asterisks indicate the site of laser severing and the yellow arrowheads indicate the position of microtubule breakage. Images were acquired using a TIRF microscope in a stream mode at a 100 ms interval. Video is sped up 90 times. Time is shown in seconds.

**Video S3, related to Figure 3. CLASP promotes formation of complete microtubules from partial protofilament assemblies.**

Microtubules were grown in the presence of Rhodamine-tubulin (magenta) and 30 nM Kin-5-GFP dimer (green) (upper left panel, corresponds to Figure 3B), in the presence of Rhodamine-tubulin (magenta), 30 nM Kin-5-GFP (green) and 30 nM TagBFP-CLASP2α (blue) (upper right panel, corresponds to Figure 3C), in the presence of Rhodamine-tubulin (magenta), 30 nM Kin-5-GFP (green) and 30 nM TagBFP-TOG2-S (blue) (bottom left panel, corresponds to Supplementary Figure S3A) or in the presence of Rhodamine-tubulin (magenta), 30 nM Kin-5-GFP (green) and 100 nM TagBFP-TOG2-S (blue) (lower-right panel, corresponds to Supplementary Figure S3B). Yellow arrowheads indicate the position of microtubule curling induced by the Kinesin-5 dimer. Images were acquired using a TIRF microscope at a 4 s interval. Video is sped up 10 times. Time is shown in seconds.

**Video S4, related to Figure 3 and Supplementary Figure S3. CLASP promotes complete repair of damaged microtubule lattices.**

Taxol-stabilized microtubules (magenta) were initially incubated in a buffer without Taxol and tubulin for 1.5 min and then transferred into a buffer with either 5 μM HiLyte 488-labeled tubulin alone (green) (upper panel, corresponds to Figure 3F), 5 μM HiLyte 488-labeled tubulin together with 30 nM mCherry-CLASP2α (middle panel, corresponds to Figure 3H) or 5 μM HiLyte 488 labeled tubulin together with 30 nM mCherry-TOG2-S (lower panel, corresponds to Supplementary Figure S3C) for 10 minutes followed by a 25% glycerol wash. Green tubulin incorporation into the lattice is indicated by numbers for the upper and middle panels and by an asterisk for the lower panel. Images were acquired using a TIRF microscope at a 2 s interval. Video is sped up 75 times. Time is shown in min.

